# Microfluidic chemostatic bioreactor for high-throughput screening and sustainable co-harvesting of biomass and biodiesel in microalgae

**DOI:** 10.1101/2022.05.06.490980

**Authors:** Guoxia Zheng, Yutong Cui, Ling Lu, Ming Guo, Xuejun Hu, Lin Wang, Shuping Yu, Shenxia Sun, Yuancheng Li, Xingcai Zhang, Yunhua Wang

## Abstract

As a renewable and sustainable source for energy, environment, and biomedical applications, microalgae and microalgal biodiesel have attracted great attention. However, their applications are confined due to the cost-efficiency of microalgal mass production. One-step strategy and continuous culturing systems could be solutions. However, current studies for optimization throughout microalgae-based biofuel production pipelines are generally derived from the batch culture process. Better tools are needed to study algal growth kinetics in continuous systems. A microfluidics chemostatic bioreactor array was presented, providing low-adhesion cultivation for algae in the gas, nutrition, and temperature (GNT) well-controlled environment with high throughput. The chip wasused to mimic the continuous culture environment of bioreactors. It allowed simultaneously studying of 8×8 different chemostatic conditions on algal growth and oil production in parallel on a 7×7 cm^2^ footprint. On-chip experiments of batch and continuous cultures of *Chlorella. sp*. were performed to study growth and lipid accumulation under different nitrogen concentrations. The results demonstrated that microalgal cultures can be regulated to grow and accumulate lipids concurrently, thus enhancing lipid productivity in one step. The developed on-chip culturing condition screening, which was more suitable for continuous bioreactor, was achieved at a half shorter time, 64-times higher throughput, and less reagent consumption. It could be used to establish chemostat cultures in continuous bioreactors which can dramatically accelerate the development of renewable and sustainable algal for CO_2_ fixation and biosynthesis and related systems for advanced sustainable energy, food, pharmacy, and agriculture with enormous social and ecological benefits.

**TEASER:** Sustainable microfluidic bioreactor for 64 times higher-throughput screening CO_2_ fixation and biomass and biodiesel production in microalgae.

## INTRODUCTION

Nowadays, the urgency of worldwide carbon-constrained environmental actions due to concerns over dangerously rising global warming and energy crisis has necessitated greater search for renewable and sustainable energy sources. Biofuel is a carbon-neutral fuel and seen as the most promising alternative petrodiesel, with particular interest in use of biodiesel and “green diesel”. [1,2] The first and second generation biodiesels have been produced from lipid feedstocks mainly derived from oilseed crops and lingocellulosic agriculture [3-5], but recently, attention has turned to utilizing microalgae for these purposes [6.7]. Microalgae, primitive plant which are called thallophytes, are one of the oldest life forms on the earth. Without any development beyond cells, algae structures still serve their primarily goals of sustainable energy conversion [7-9] and biomedical applications [10, 11]. They could convert the green house gas and sun light into chemical energy through their conventional physiological pathway, part of which is what we used as biodiesel (Fig.1a). Many algal strains are able to accumulate in certain conditions, high amounts of fatty acids in up to 20-50% of their dry weight [12]. Additionally, microalgae have faster growth rate, less freshwater demand for cultivation, no or minimal competition with food supply, land usage and associated environmental impacts since microalgae could live in very harsh environment, such as saline/brackish water/coastal seawater on non-arable land [6-8]. Noticeably, microalgae have superior CO_2_ fixation capacity, minimizing CO_2_ footprint, especially their growth could be coupled to the direct bio-fixation of waste flue gas [8, 13]. Despite these tremendous potentials, the concept and feasibility of microalgal biodiesel have been discussed extensively. The principal contradiction is that the production cost of current algae derived biofuels and bioproducts remains well above economic viability. Significant improvements are still required throughout all steps of the microalgae-based biofuel/bioproduct production pipeline [6, 7, 12].

In general, neutral lipids are in the form of Triacylglycerols (TAGs), the secondary metabolites of microalgae, which are synthesized via transesterification process in algae by use of batch type reactor [14]. This process begins at the end of exponential growth phase and executes rapidly in the cellular stationary phase. In these phases, nutrients are almost depleted, resulting in non-homogeneous conditions and subsequent low efficiency in cell growth. Although oil content per cell could be intensively accumulated in such stage, the decreased total amount of biodiesel production could occur [14, 15]. To overcome this contradiction, “two-stage cultivation” strategies in bioreactor have been explored to enhance microalgal lipid production. In such strategies, Microalgae first grow rapidly in growth-optimized batch culture, and then are transferred to culturing conditions in which physical or chemical inducible factors, such as light intensity, temperature, nutrition, culture pH and CO_2_, *etc*, are adjusted to promote lipid accumulation at the cost of cell growth[16]. However, detailed study of the stress responses of lipid-rich microalgae in continuous culture may provide a mutual benefit for the production of biomass and lipids in one step, a strategy that could be more suitable for mass production. Unfortunately, it is an open question how microalgal species will respond in continuous culture. Previous studies demonstrate that the growth of some algal species (e.g. *Chlorella pyrenoidosa, Chlorella vulgaris*) were seriously hampered under high nitrogen stress, but lipid accumulation were induced before the growth limitation. While others (e.g. *Neochloris oleoabundans, Choricystis minor*) produce lipid in relatively high nitrogen stress conditions, their growth were not significantly limited [17-19]. These features may or may not be interspecies-specific characteristics, have inspired great deal of explorations for the threshold values of nutrition concentration, in order to establish a continuous culture system for win-win situation of biomass and lipids [18-21]. However, the available bioreactors are straightforward to use. They are not designed for efficient system parameter optimization. Current microalgal studies are usually conducted by culturing the organism in lab-scale flasks and multi-well plates [22, 23]. These culture systems have made significant contributions to the understanding of basic algal biology, selecting the best strains for the production of biochemical, and understanding the effects of various culture factors (e.g., light intensity, temperature, nutrient concentration, CO_2_, pH) on algal growth and oil production [22, 23]. Unfortunately, these systems are all batch process systems, suffering from time-dependent nutrient depleting. This non-homogeneous culture state is substantially different form that in chemostatic bioreactor with continuous nutrient supply. In addition, a heterogeneous state makes it difficult to analyze cellular properties due to the many-conditions-to-one-phenotype features [14, 24, 25]. Better tools are needed to study algal growth kinetics thus can accelerate our understanding of such managed engineered continuous systems (Fig.1a).

Microfluidic lab-on-a-chip systems, with their capability to precisely control, monitor and manipulate samples at the nano to pico-liter scales, as well as to integrate various steps in a particular biological assay, are ideal for creating a high throughput screening platform [26, 27]. A few microfluidic systems examine microalgal motions, lipid production, density changes, or growth kinetics [28-30]. However, these systems focus on providing batch culture condition. Some systems that provide semi-continuous or continuous cultures relying on endogenous valve switching, micro-droplet fusion/division, or film diffusion, are suffering from the difficulties in fabrication or operation, and therefore, not suitable for high-throughput screening applications[31-33]. Another factor hampering the mcirofluidic performance [34, 35] is the unwanted adsorption on hydrophobic (polydimethylsiloxane) PDMS surface, the most widely used biocompatible material in fabricating cell-based microfluidic devices. Unfortunately, most probes and markers (i.e. BODIPY and Nile red) used in algae lipid analysis have a hydrophobic properties, so they readily absorb on PDMS surface, thus rendering on-chip fluorescence lipid imaging impossible. Furthermore, learning from our experimental experiences, although algae are almost floating-cells, even motile cells, increasing non-specific adsorption on culturing chamber surface (glass, silicon wafer, PDMS, polyster film, *etc*.), mightily resulting from increasing probability of collision in micro-scale, has occurred in long-term culturing. It will decrease the cellular surface available for nutrient uptake while maintaining the cells’ physiological properties.

The high-throughput bioreactor array presented here provided low-adhesion cultivation for algae under an extremely well-controlled chemostat environment at high throughput. As shown in Fig 1b, the array is composed of 8 identical units which consist individually of an upstream dynamic controllable nutrient supply base and a downstream Polytetrafluoroethylene (PTFE)-coating cell culture array. The cell culture array is connected with nutrient supply channels by micro-diffusers which are impassable for cells, but allow for effective molecule diffusion, and thus for continuous nutrient feed and preventing accumulation of excess metabolites. This bioreactor array is capable of simultaneously studying the effect of 8×8 different chemostatic conditions on algal growth and oil production. Coupled with arrays of miniaturized microalgal culture chambers, 64 independent bioreactor experiments could be conducted in parallel on a 7 × 7 cm^2^ footprint. Besides chip, we developed a mating system that allows gas, nutrition and temperature (GNT) controlled culture as well as in situ determination (Fig. 1a). These controlled conditions plus the core chip were integrated and used to mimic the continuous culture environment of bioreactors, where the CO_2_ fixation and biosynthesis of microalgae happen. Our GNT-chip system enable the simulate chemostatic bioreactor in situ studies of diverse biological processes in ways that are not possible to be accomplished by using conventional algae study bench-tops, benefiting algae biodiesel/byproduct studies. On-chip experiments of batch and continuous feed-batch cultures of *Chlorella. sp*. (chl-1, KLEMB, IOCAS) were performed to study its responses, in terms of growth and lipid accumulation, to different nitrogen concentrations. The results were used to establish chemostat cultures in bioreactor with different dilution rates, in order to verify the functional mcirofluidics and also the assumption that microalgal cultures can be regulated to grow and accumulate lipids concurrently, thus enhancing lipid productivity.

**Figure 1.**
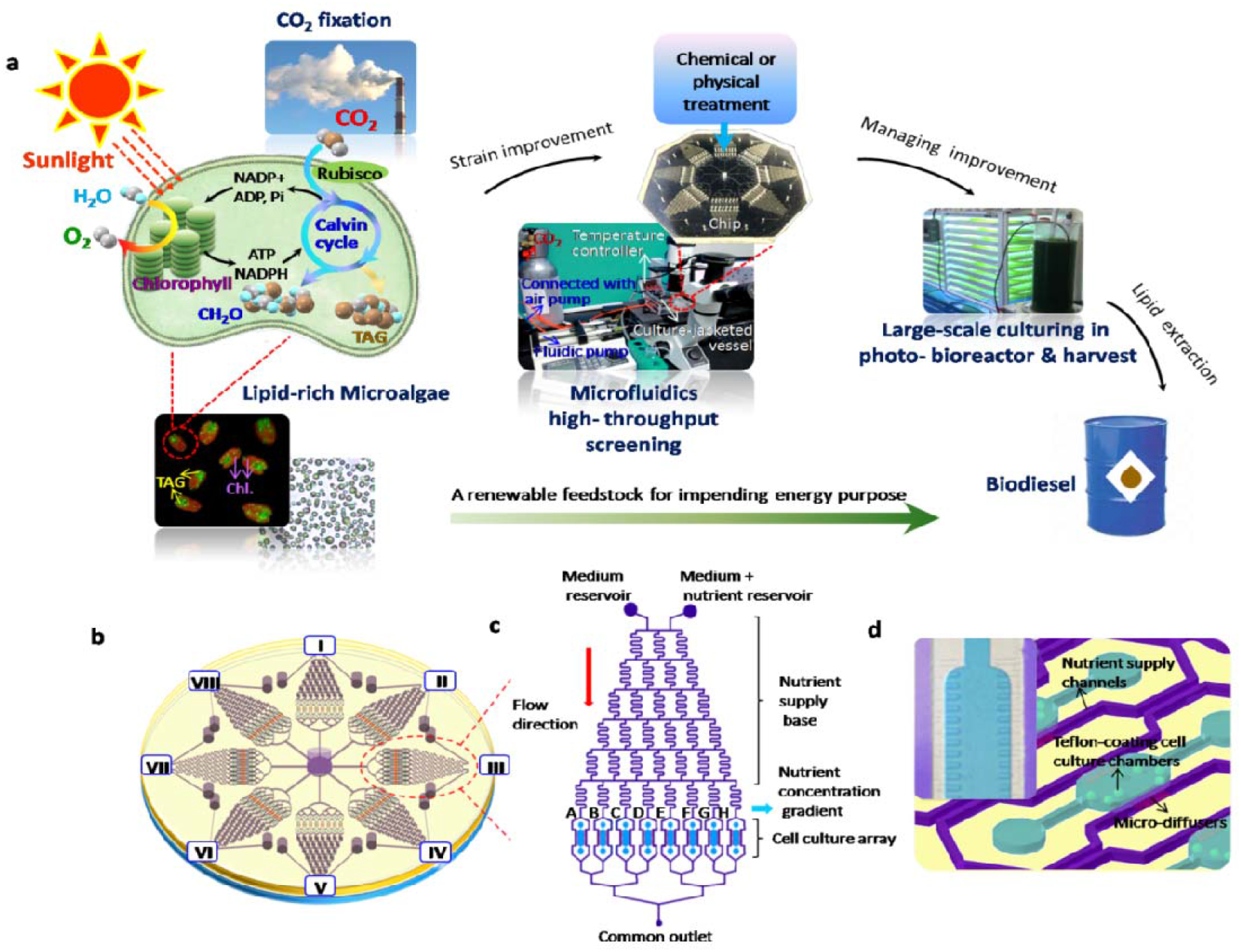
Microfluidic-based continuous culturing system toward high-throughput screening of co-harvesting biomass and biodiesel in microalgae. (a) Schematic illustration of all steps of the microalgae-based biofuel production pipeline and how the developed GNT- Microfluidic chemostatic bioreactor system work. Algae cell structure serves goals of biodiesel production through the photosynthetic carbon fixation pathways-the Calvin-Benson cycle. (b) Microfluidic designed for simultaneously studying the 8×8 different chemostatic conditions on algal growth and oil production; (c) One functional unit of the microfluidic; (d) Structure of nutrient supply channel, micro-diffuser channel and micro-culturing chamber.

## RESULT AND DISCUSSION

### GNT-Chip system design and characterization

Usually microalgae growth, i.e. biomass accumulation and TAG synthesis do not run in parallel through cultivation, which necessarily lowers overall lipid productivity. In-depth analyzing the stress response of lipid-rich microalgae in continuous culture may provide a co-harvesting of biomass and lipids in one step that a strategy will benefit large-scale algae-based biodiesel production.[18-20] However, due to the methodological limitation, the available algae study bench-top apparatus can neither provide homogenous culturing conditions, which is substantially different form that in chemostatic bioreactor, nor yield a well-controlled high-throughput operation. Microfluidics, also known as “lab-on-a-chip systems”, an emerging technology which represents a revolution in laboratory experimentation, bringing the benefits of integration, miniaturization, and automation, can offer attractive alternatives for current cell culture and handling technologies to advance the current state of microalgal biofuel/bioproduct research. [26, 27]

Herein, we developed a GNT-chip system to actively simulate the chemostatic bioreactor to high-throughput screening in situ the stress applied to microalgae in gas, nutrition and temperature (GNT) controlled continuous culture. The detailed construction process consists of encapsulating a micropatterned microfluidic chip in a PMMA matting holder with a temperature controller as well as a gas controller and a fluid pump (Fig. 1a). Parallel PDMS microchambers with high thermal conductivity and air permeability guaranteed that each algae cell was cultured and analyzed under controlled constant temperature and CO2 atmosphere. The core chip is a high-throughput microfluidic bioreactor array capable of investigating the effects of 64 different nutrition conditions on microalgal growth and oil production. The key feature of our array design is an aesthetic spectacle that eight identical network structures of central symmetry are precisely patterned in a 7 × 7 cm^2^ footprint PDMS piece (Fig.1b). For each unit, it is an integration of downstream PTFE- coating cell culture chambers flanked by two side feeding channels which were connected with an upstream nutrient supply base (Fig.1c). Dynamic controllable and continuous feeding of nutrients to each culture compartment where the walls covered with “no-sticky” paint coat allowed long-term culturing and low signal-to-noise ratio (SNR) in visual analysis (Fig.1d). This platform also overcame the limitations of conventional culture systems by applying homogenous conditions to all microalgae in confining but no trapping sites and implementing high-throughput screening capabilities.

In each nutrient supply base, as depicted in figure 1c, a series of winding channel networks utilized inherent laminar flow and diffusive mixing to generate a successive combinatorial mixture of two inputs (seawater and nutrition solution). The exiting streams containing nutrient concentration gradients feed into a downstream array of diffusible chambers. To assess this concentration gradient generating behavior, 0% rhodamine 123 (i.e., Diluted Water) and 100% rhodamine 123 solutions were introduced separately into the microfluidic chip. It was observed that concentrations at the eight outlets in each supply base were all linearly generated (Fig. 2a). The correlation factors between the theoretical estimations and the experimental data for the eight units are 0.984-0.995 [36], verifying the feasibility of this chip to generate multiple well-defined concentration gradients simultaneously (Fig. 2b). It will well-suitable for the detailed screening of the stress response of lipid-rich microalgae in continuous culture by properly regulating the nitrate concentration (e.g., severe stress or moderate stress). Additionally, large-scale dose-response tests can be performed in a simple way as the culturing conditions are flexible using supply base and can be optimized by changing either the category of chemicals (e.g. nitrate, phosphate, urea, glues, pH, HCO_3_^-1^, etc.) or the concentration of the chemicals.

**Figure 2.**
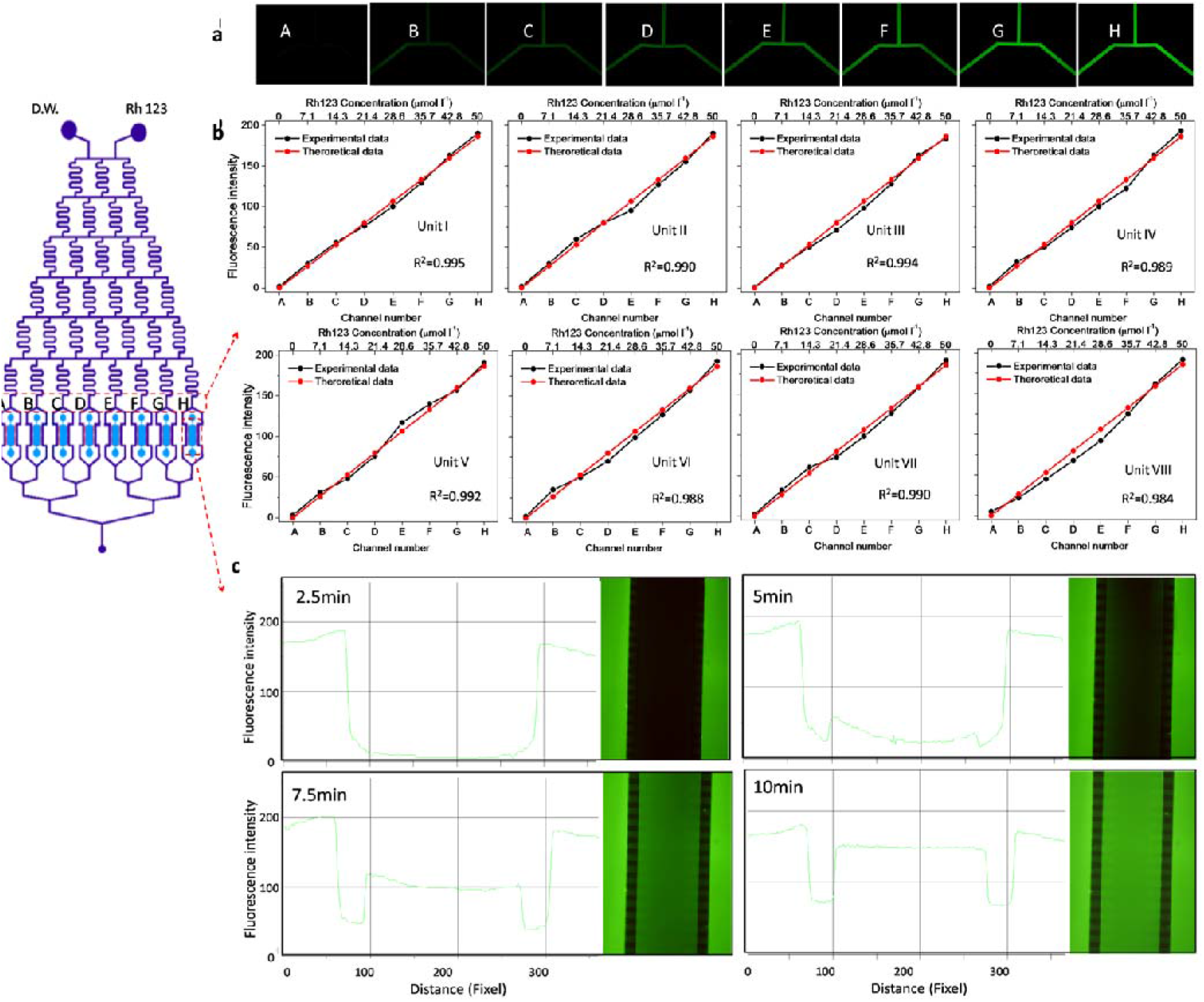
Characterizations of on-chip nutrient supplies. (a) A series of solutions with different concentrations of Rh 123 were formed at the eight outlets of represented nutrient supply base. (b) The intensities of Rh123 at eight outlets of each nutrient supply base by inverted fluorescence microscope were quantified and compared with the theoretical value. (c) The Rh123 diffusion in the chamber versus time.

The downstream functional area is an array of parallel feeding channels with continuous flow through of fresh medium and parallel rows of cell culture chambers between the channels. This module is designed to prevent any active flow through the chambers and possible perturbation of cell position, motion or intercellular interaction. The branching from a single channel connected with nutrient supply base into the array of 16 parallel nutrient supply channels is binary and symmetric. This arrangement leads to equal flow rates through the nutrient supply channels, highly balanced pressures at binary channels on opposite sides of the chambers and thus negligible flow through the chambers. To one-dimensional approximation [37], the time required to establish a steady condition in chambers is t ≈ l^2^/2D, with D = ∼10^−10^ m^2^ s^−1^ and l = 300 μm, which is 7.5 min. Experimental results were further validated against results from this estimated time. Figure 2c depicted the diffusion of Rh 123 into the culture chamber. Insert photos were processed to obtain the changes in fluorescent intensity across the chamber at 2.5min, 5min, 7.5min and 10min time point, respectively. Fluorescent intensities were calculated from one of the branching parallel channel to another for all pixels (distance) across the length of the chamber. At 10 min time point, the intensity profile across the chamber presented as a nearly straight line. It implied that nutrition condition inside the culture chambers could be homogenous and identical with that in the nutrient supply channels within 10 minutes. It will be valuable in fundamental and applied chemostatic bioreactor simulation and continuous culture research.

Besides supplying chemostatic cultivation conditions to all microalgae, our culture array design provides cell confinement for both motile and non-motile cells because microchambers are not directly interconnected with feeding channels, but are bridged by micro-diffusers of depth of ∼3μm, which are impassable for cells, but allow effective molecule diffusion (Fig.1d). This is important because many microalgae are highly motile [38]. Confine ability of diffusible chambers was assessed by examining the cell concentration of motile microalgae in plateau phase over the 24 h period. Figure s1 shows cell concentrations of *Platymonas subcordiformis* and *Platymonas helgolandica var. tsingtaoensis* at the 0 and 24 h time points, and it is seen that cell concentrations for these two microalgae remained statistically equivalent during the duration of the experiment (*P*>0.05), verifying that mobile microalgae can be successfully confined in the culturing chambers.

To eliminate the adsorption of small molecular fluorescence probe or floating cells, the surface of culturing chambers were coated with PTFE(brand name Teflon®), which is known to have both oleophobic and hydrophobic properties whose treatment will serve to confer a degree of small molecule absorption resistance to PDMS. To assess this resistance, we tested that a Teflon-AF1600 coating resisted small molecule absorption of Rh 123 (MW=380.824). Figure 3 shows the results for unmodified and PTFE-coated PDMS microchannels. After 3 h of exposure, the no-coating PDMS channel exhibited significant adsorption of Rh123 through the side walls of the microchannel (Fig. 3a, b), resulting in very strong background fluorescence signal even after rinsing (Fig. 3c). The PTFE-coating microchannel had no absorption of the Rh123 physical geometry. The significantly enhanced non-smearing effect for rhodamine dye indicates that we successfully achieved a small molecule resistant PDMS Teflon coating.

**Figure 3.**
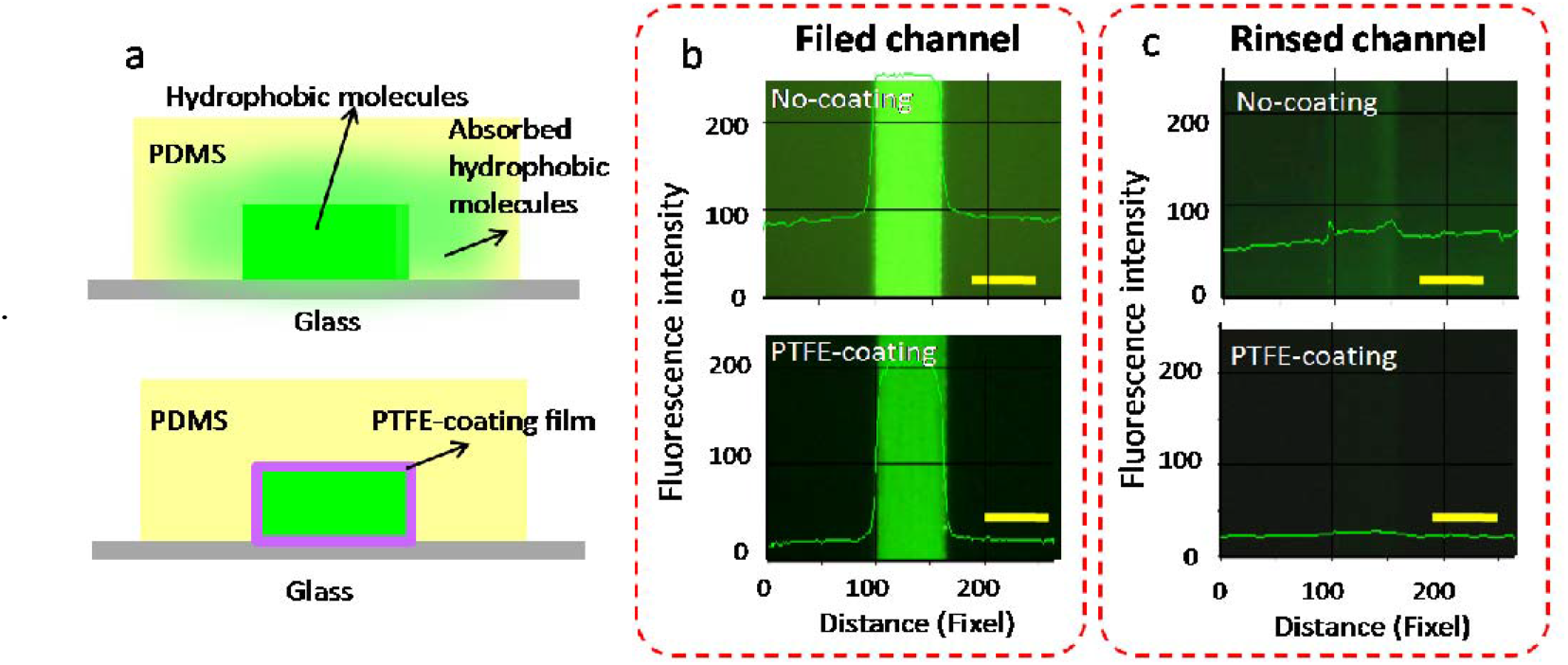
(a) The resisting of PTFE-coating PDMS towards small molecule absorption Fluorescence images of (b) Rh 123 filed channels and (c) rinsed channels showing strongly Rh123 adsorption on no-coating PDMS, but greatly eliminated adsorption on PTFE-coating PDMS.

The elimination of hydrophobic molecule adsorption is also demonstrated by use of BODIPY493/503, a well-known hydrophobic fluorescence probe, which was widely used to visualize intracellular lipid droplets due to its high sensitivity and ease of use [30]. As a demonstration example, *Chlorella. sp*. (chl-1, KLEMB, IOCAS) were loaded into the PDMS bioreactor with micro-diffusers while flowing 10 μg ml^-1^ BODIPY in feeding channels for staining. The cells were confined within chambers and incubated for 20min, and then flowing D.W. in feeding channels for rinsing residual dye within chambers before observation. As seen in figure 4a, it is difficult to visualize intracellular lipids in *Chlorella. Sp* cells due to the strong background fluorescence signals resulting from unable rinsed PDMS absorption within unmodified chambers. However, in PTFE-coating chambers, their lipid signals were clearly distinguishable with generally dark background. The lipid signal intensities among different algae species in chambers were further observed. As seen in figure 4b, the lipid bodies in all four algal species exhibited clearly bright green fluorescence, although their intensities varied markedly. We visualized the green fluorescence and calculated them using image J software. The off-chip comparative experiments were performed by total lipid extraction method [17]. As illustrated in figure 4c and d, both on-chip and off-chip lipid accumulation measurements showed inter-species markedly distinguishable where *Chlorella sp*. exhibited higher lipid content than other three and can be the candidate for biodiesel production (*p*<0.05). These demonstrations strongly supported that the surface modified micro-bioreactor reported in this paper allows to detect optical fluorescent signals even distinguish interspecies heterogeneous lipid accumulation by inhibiting the absorption of hydrophobic molecules such as BIODIPY on PDMS.

**Figure 4.**
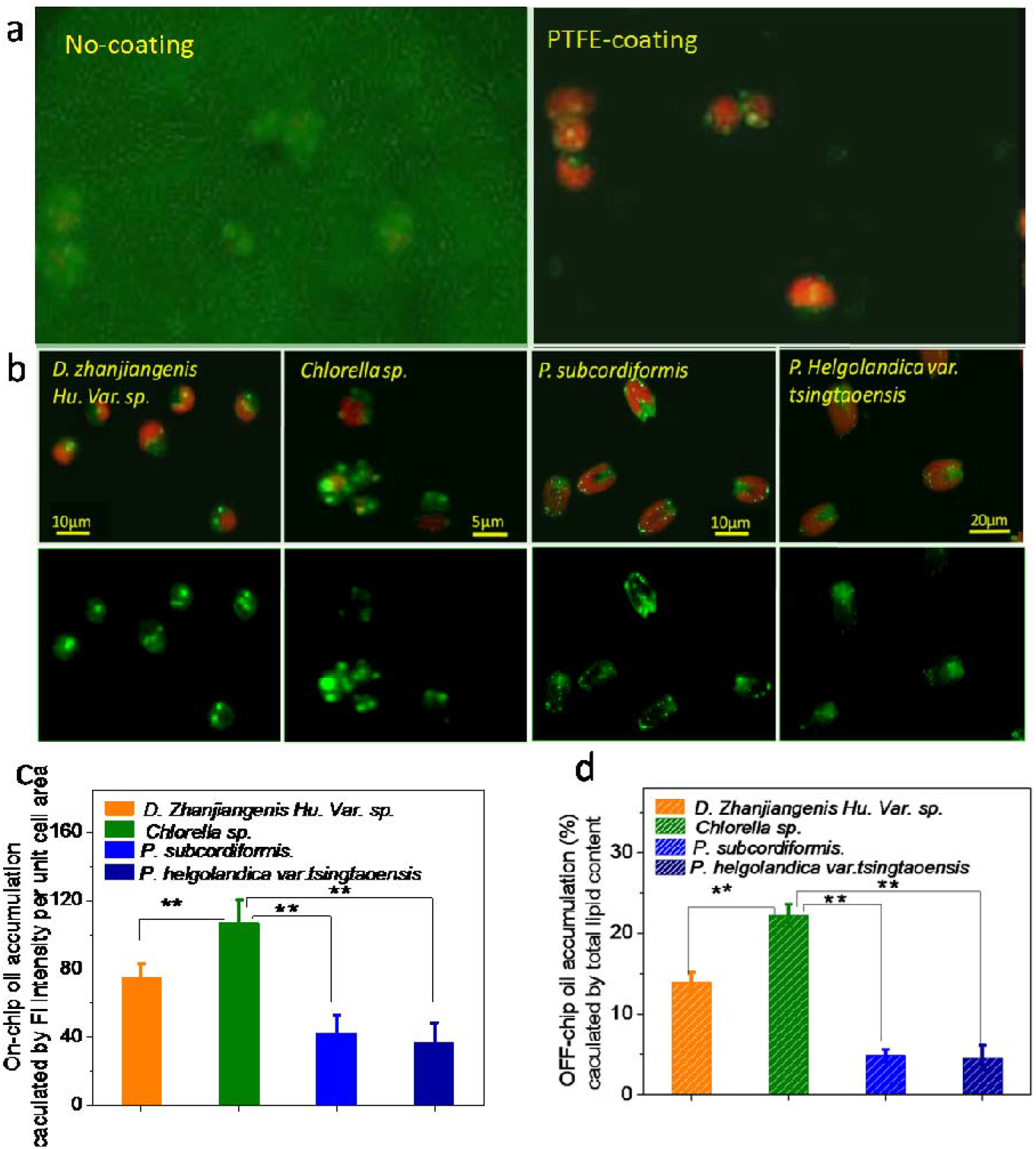
Micrographs showing the fluorescence of lipid droplets stained with BODIPY (green) in unmodified and PTFE coating PDMS microchambers. (a) Observation of intracellular lipid in the unmodified device is almost impossible due to the interference of adsorbed fluorescent probe. However, in PTFE-coating chambers, lipid signals were clearly distinguishable with generally dark background that (b) interspecies comparison of lipid productivity in microalgae was successfully performed. (c) Average fluorescence intensity of BIODY stained four strains of algae were analyzed after 5 days’ culturing. (d) Off-chip measurements of oil accumulation for 5 days’ culture.

Furthermore, to assess the resistance of cell retention on modified surface, micro-chambers with a “chessboard” patterned PTFE-coating glass surface were prepared (Fig. 5a). *Giardia trophozoites*, a sedentary single-cell parasite, as a demonstration example, were loaded into the modified chambers. After 24 h static incubation, the number of *Giardia* on PTFE-coating chessboards is generally similar to that on naked glass chessboard. (Fig.5b). And then the cells within chambers were treated with flow-induced shear stress by connected with a fluidic pump. After the shear stress challenge, the percentages of *Giardia* remaining attached on both surfaces were counted. As seen in figure 5c-d, the cell percentage remaining attached on PTEE-coating surface is significantly lower than that on glass surface. Obviously, non-specific interactions between cell and solid-phase surface that are derived from contact force (a combined electrostatic and van der Waals force), were generally un-sustained on Teflon surface due to its chemical structure. Only 8% *Giardia* kept stationary on PTFE-coating surface under high flow stress. It attributed to the negative press derived from ventral disk, a special physiological structure of *Giardia*. The bioattaching resistance property of Teflon was further tested based on a method of digital microfluidic (DMF) actuation. A long-term static incubation (15 days) of microalgae *P. subcordiformi* within droplet among two parallel PTFE-coating electrode plates was performed and then the droplet was actuated. As shown in figure 5f and the supplementary video, almost all algal cells remained in suspension and moved following the movement of droplet. Generally none (∼< 9‰) cell attaching occurred after a long-term culturing. These results indicate that PTFE-coating properties can substantially inhibit unwanted algae attachment and thus maintaining the cellular contact area available for nutrition uptake. It will be valuable in fundamental and applied microalgae-based research.

**Figure 5.**
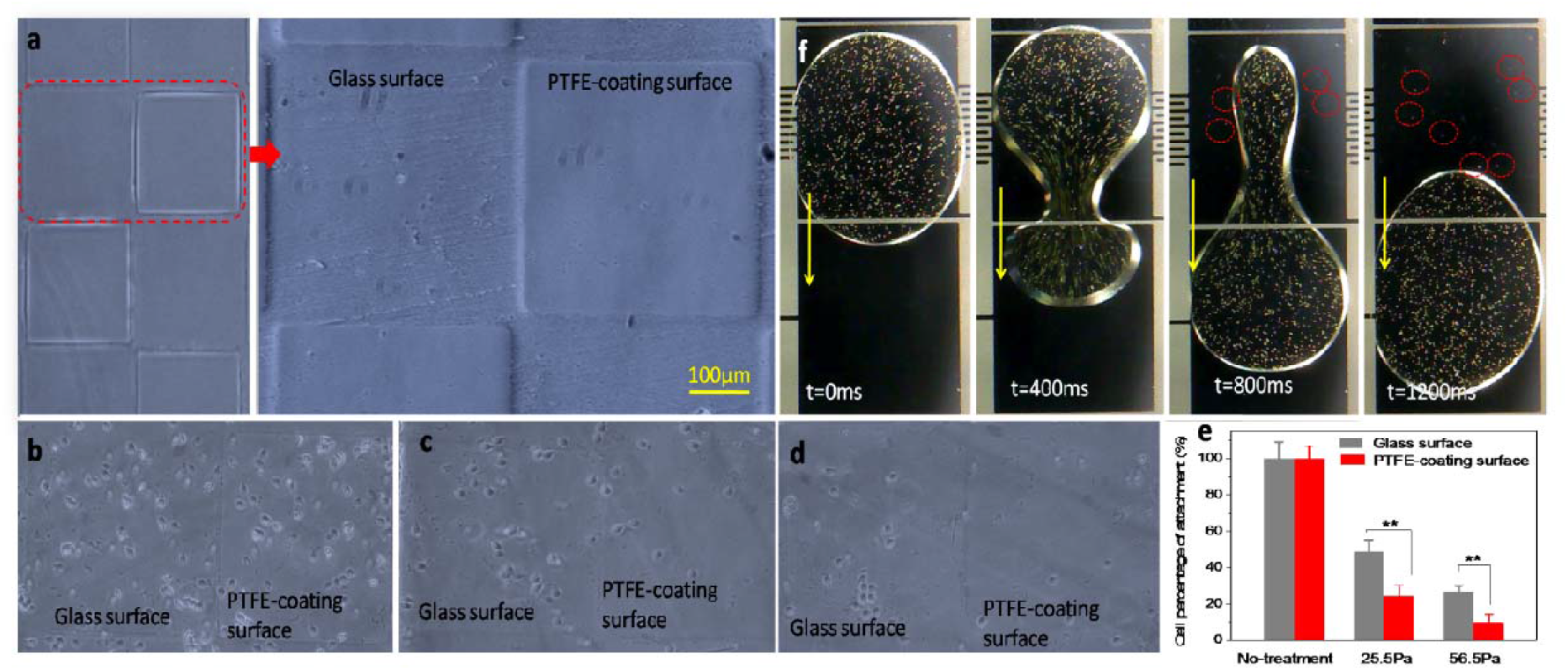
Resistance of cell retention on PTFE modified surface. (a) “Chessboard” patterned PTFE-coating glass surface. (b) Attachment of *Giardia* trophozoites on “chessboard” patterned PTFE-coating glass surface. After 24 h static incubation, the number of Giardia on PTFE-coating chessboards is generally similar to that on naked glass chessboards; (c) Challenged by a shear stress of 25.5 Pa; (d) Challenged by a shear stress of 56.5 Pa; (e) The *Giardia* cell percentage remaining attached on PTEE-coating surface is significantly lower than that on glass surface. Only 8% *Giardia* kept stationary on PTFE-coating surface under shear stress of 56.5 Pa; (f) Incubation of microalgae *P. subcordiformi* within droplets among two parallel PTFE-coating electrode plates. Almost all algal cells remained in suspension and moved following the movement of droplet.

### Microalgal growth kinetics in the array microbioreactor

Using the microfluidic device, we first studied growth kinetics of the microalga marine *Chlorella sp*. in both batch mode and continuous feeding mode. We once and continuously flow the commonly used f/2 media along the feeding channels, respectively. Fluorescent images of the microbioreactor array were taken every 24 hours. Cell growth with time within each microchamber is clearly shown in figure 6a. Using the fluorescence images, we counted the cell numbers with image J. Meanwhile, the data were also obtained by a flask scale in order to set up the correlation between the data produced in a picoliter or nanoliter volume microchamber and a flask scale, which made the on-chip data meaningful to the researchers who ordinarily operate the microalgal culture in a flask scale. The resulted growth curves for two on-chip culturing modes and flask scale all demonstrated a lag phase and exponential growth phase followed by a stationary phase (Fig. 6b). The microalgal growth curves cultivated at batch mode on the microfluidic devices and on the flask scale agreed well. Their growth characteristics, such as the length of the lag phase, the specific growth rate, and the maximum cell concentration, were found comparative. It was consistent with the results with the same media done in our lab and others [8, 39-42]. These comparative analysis results indicated that batch growth on the microfluidic device could be associated with scaling the system to the flask scale.

**Figure 6.**
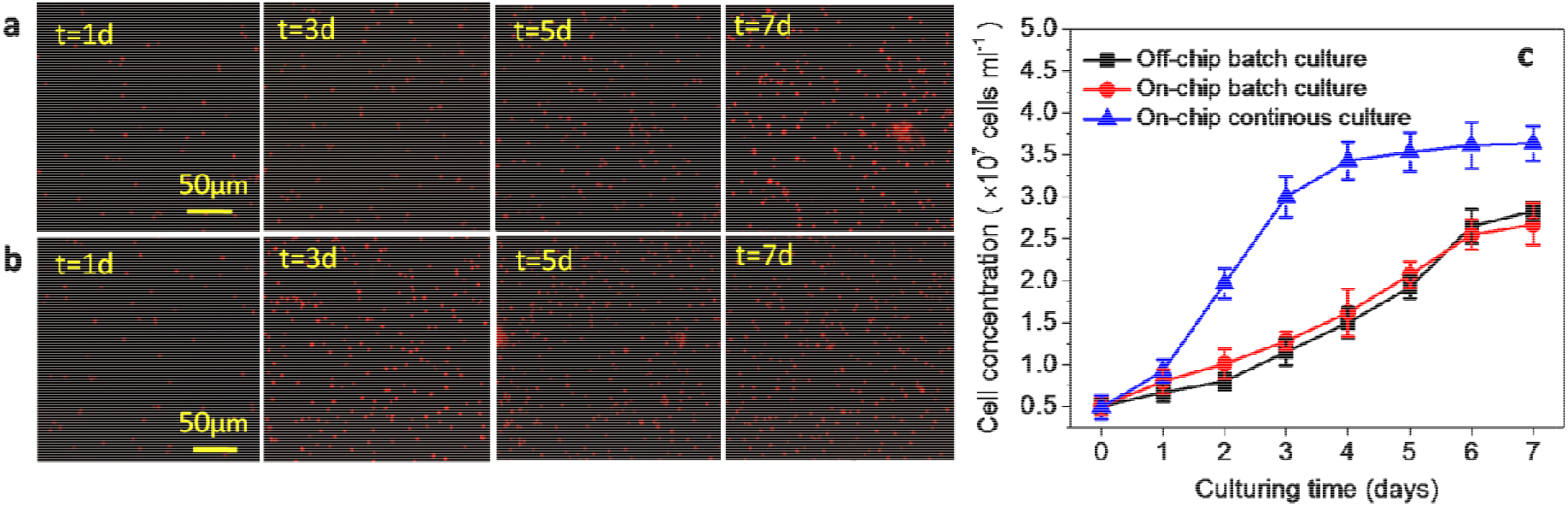
Growth kinetics of the microalga marine Chlorella sp. (a) on-chip batch mode; (b) on-chip continuous feeding mode; (c) The specific growth rates were calculated from slopes of the growth curves in the exponential region : 0.582 ± 0.13 (on-chip continuous mode) day^−1^, 0.265 ± 0.09 (on-chip batch mode) and 0.270 ± 0.07 (off-chip batch mode, flask scale) day^−1^. The maximum cell concentrations after 7d-incubation period were found to 3.56×107, 2.61×10^7^ and 2.69×10^7^ cells ml^-1^, respectively.

The microalgal growth characteristics at on-chip continuous mode were found to vary significantly with that at batch mode on the microfluidic device and on the flask scale. The slopes of the growth curves in the exponential region provided us the specific growth rates: 0.582 ± 0.13 (on-chip continuous mode) day^−1^, 0.265 ± 0.09 (on-chip batch mode) and 0.270 ± 0.07 (flask scale) day^−1^. The maximum cell concentrations after 7d-incubation period were found to be 3.56×10^7^, 2.61×10^7^ and 2.69×10^7^ cells ml^-1^, respectively. These results were caused by the difference in the nutrient feeding method. The nutrients were supplied continuously on the continuous-feeding microfluidic chambers, whereas nutrients were supplied by once under the microfluidic and flask-scale cultivation conditions, resulting in a rapid nutrient depletion and subsequent low efficiency in cell growth. Noticeably, the elapsed times required to reach the stationary phase at continuous mode and two-stage batch were 3 and 6 days, respectively. In the stationary phase, nutrients are almost depleted, algal carbohydrate accumulation declined, while the fatty acid accumulated intensively. Thus the time required for screening of culture conditions that induce high growth rate and oil production on a continuous-feeding chip was much shorter than that required on the flask scale.

### Nitrate-stress cell growth and lipid accumulation in on-chip batch culture

Nitrogen starvation has frequently been reported to induce lipid accumulation in microalgal cells [17-21]. To illustrate the growth and lipid responses of *Chlorella sp*. (chl-1, KLEMB, IOCAS) to variant nitrate levels, a series of batch cultures were performed on chip at different initial nitrate concentrations. After 7 days’ culturing, on-chip in situ profiling of the nitrate starvation triggered growth and lipid contents were shown in figure 7. As shown in the enlarged fluorescence images, N (NaNO_3_) fertilizers have a great influence on the biomass and lipid content of *Chlorella sp*. (Fig.7a and b). It was found that the increase of nitrogen concentrations would greatly promote the growth of cells to produce more biomass. However, accumulated lipid bodies exhibited highly increasing fluorescence signals among cells with a decrease in nutrition concentration. The lowest cell density but the clear/large cytosolic lipid bodies are seen in 0.22mM concentration of nitrate. The resulting growths are shown in figure 7c (Blue line). These results showed that specific growth rate of *Chlorella sp*. (chl-1, KLEMB, IOCAS) increased almost linearly with increasing initial nitrate from 0.22 mM to 0.88 mM. Further increases in the nitrate concentration will not increase biomass accumulation. The lipid content, demonstrated by fluorescence intensity per unit cell area, however, exhibited different trends in response to increasing nitrate concentration (Fig. 7c, Red line). Specifically, the lipid content first slightly fluctuated from 150 units to 120 units, and then to 100 unit, as the initial nitrate concentration was increased from 0.22 mM to 1.76 mM. Further increases in the nitrate concentration will decrease lip accumulation significantly. These results showed that cell growth and lipid accumulation did not always change in parallel, and moreover, they suggested relatively high lipid content could be achieved without the expense of algal growth. As shown in figure 7c, decreasing the nitrate concentration from 1.76 mM to 0.44 mM significantly increased the lipid content (*p < 0*.*05*), while the biomass was not severely affected (*p > 0*.*05*).

**Figure 7.**
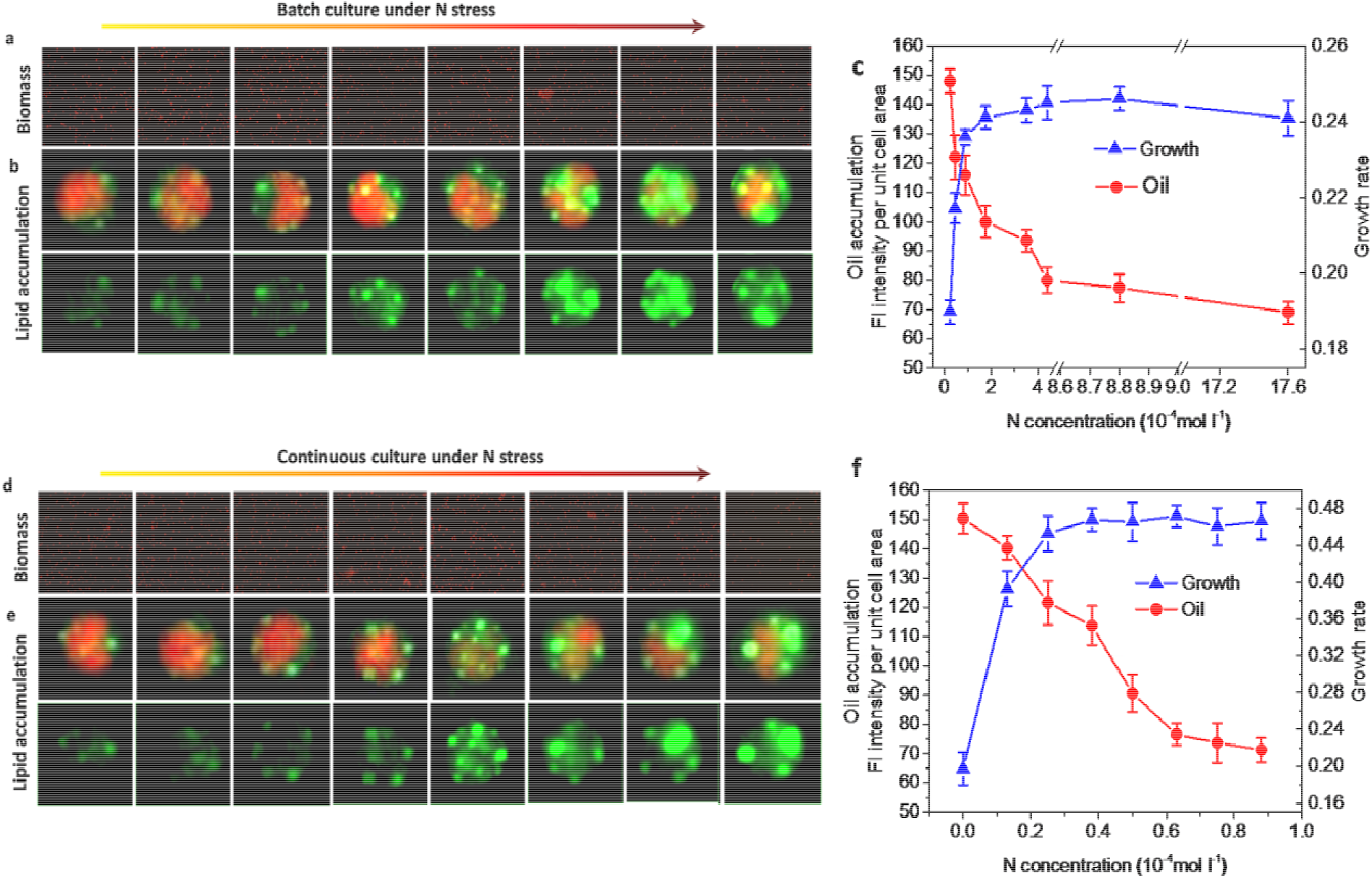
On-chip in situ profiling of the N stressor triggered cell growth and lipid accumulation. Enlarged fluorescence images of (a) algae cells populations and (b) cytosolic lipid bodies in batch culture. (c) The growth and oil accumulation measurements of *Chlorella sp*. in batch culture. Enlarged fluorescence images of (d) algae cells populations and (e) cytosolic lipid bodies in continuous culture. (f) The growth and oil accumulation measurements of *Chlorella sp*. in continous culture.

### Effects of continuous nitrate feeding on cell growth and lipid accumulation

To confirm that there is a culture status under which *Chlorella sp*. (chl-1, KLEMB, IOCAS) moderately accumulates lipids in growing cells, 4 days’ continuous culture was conducted with nitrate concentrations of 0.00, 0.13, 0.25, 0.38, 0.50, 0.63, 0.75 and 0.88 mM. A similar response of biomass and lipid content to varying nitrate was observed (Fig. 7d-f). In particular, the cultures with nitrate concentration of 0.25 and 0.38 mM almost reached maximal biomass density of about 3.2×10^7^ cell ml^-1^when nitrate was fed continuously, and all of them showed fairly high lipid content (113–121units). These results suggested that microalgal growth and lipid accumulation could both be optimized by using continual feeding of small amounts of nitrate. This on-chip screening optimum status of 0.18 mM nitrate concentration for concurrent microalgae growth and lipid accumulation can be further used in large-scale bioreactor. To verify the functional mcirofluidics and on-chip results, the chemostat cultures of *Chlorella sp*. (chl-1, KLEMB, IOCAS) in bioreactor were investigated as follows. Steady-state studies of *Chlorella sp*. (chl-1, KLEMB, IOCAS) were carried out using a chemostat system with constant medium composition (f/2 with 0.18 mM nitrate), with dilution rate of 1.0 d^-1^, respectively. Batch culture of *Chlorella sp*. chl-1 was carried out in the same photobioreactor used for the continuous cultures in order to supply basic data for comparing chemostat and batch cultures. 7-days biomass productivity, lipid content and lipid productivity were determined and compared in Table s1. The chemostat culture with dilution rate of 1.0 d^-1^ had twice the biomass productivity of the batch culture, while the lipid content was somewhat lower than that in the batch culture. Due to the sustained algal growth and moderate lipid accumulation, the lipid productivity of the chemostat (64.89 mg l^-1^ d^-1^) was significantly higher than that of the batch culture (47.04 mg l^-1^ d^-1^). Hence, in comparison to batch culture, lipid productivity can be significantly enhanced by continuous cultivation of oleaginous microalgae with proper specific nitrate input rate. This new strategy has a great advantage over batch culture in both biomass productivity and lipid productivity. When it is applied to mass culture of oleaginous microalgae, the cost of microalgal lipid production could be reduced significantly, thus promoting the commercialization of microalgal biodiesel. This work is also an advancement in the field of sustainable Nature/bio-derived materials [43-48]. It can dramatically accelerate the development of renewable and sustainable algal for CO_2_ fixation and biosynthesis and related systems for advanced sustainable energy, food, pharmacy, and agriculture with enormous social and ecological benefits [9, 10, 26, 27, 49-55].

## Conclusion

A high-throughput microfluidic microalgae bioreactor array was developed to investigate growth and oil production of microalgae under well controlled nutrient conditions. The unique features of the system are the PTFE-coating and diffusion-based microhabitat array format suitable for studies of microalgal cell (both motile and non-motile) with more sensitive optical signals, which represents the advancement in the miniaturization of the continuous cell culture system. *Chlorella sp*. chl-1 colonies were successfully characterized using the developed platform. The results showed that both microalgal growth and lipid accumulation could be optimized by using continual feeding of small amounts of nitrate. This identified nutrient condition was further used in a large-scale bioreactor. Enhanced lipid productivity demonstrated the feasibility of one-step production of microalgal lipids by continuous culture, which provides a new strategy in development of microalgal biodiesel.

The developed on-chip culturing condition screening, which was differ from conventional batch cultures and was more suitable for continuous bioreactor, was achieved at a half shorter time, 64 times higher throughput and less reagent consumption. We expect that this platform will serve as a powerful tool to investigate how algal growth and oil production are influenced by various growth conditions against algal strains of interest, at significantly lower cost and shorter time, which can dramatically accelerate the development of renewable and sustainable algal for CO_2_ fixation and biosynthesis and related systems for advanced sustainable energy, food, pharmacy, and agriculture with enormous social and ecological benefits.

### EXPERIMENTAL PROCEDURES

#### Microfluidic chip design and fabrication

The schematic of the microfluidic chip for high-throughput bioreactor array (size: 7 × 7 cm^2^) is shown in Fig. 1b-d. The chip consists of eight identical structure units and each unit contains of upstream dynamic controllable nutrient supply base and a downstream Teflon-coating cell culture array. Eight units share the central removable outlet connected with a tube. The nutrient supply base utilizes a series of diffusive mixing channel networks through which different dilutions of chemicals are automatically generated at outlets from two fluid inlets. By flowing seawater and nitrate solution through each inlet, the 8-outlet nutrient supply base produces 8 different concentrations of nitrate solution into downstream nutrient supply channels. These nutrient supply channels circled round downstream cell culture chambers (depth: 100μm, width: 500μm, length: 2mm) with a 50μm gap. They are only connected by micro-diffusers (depth: ∼3μm, width: 30μm), which are impassable for cells, but allow for effective molecule diffusion, and thus for screening microalgae against 8×8 different continuous feed conditions in parallel.

The micro-bioreactor array was prepared using PDMS by soft lithography. Photolithography using SU-8 negative photoresist (Microchem, USA) was used to generate a master mold on a Si wafer (Unisill Wafer, Korea), which was firstly patterned to form a 3 μm-thickness for defining the height of micro-diffusers that could be secondarily patterned to form a 100 μm-thickness in a subsequent photolithographic step for channels and chambers. A PDMS piece was then fabricated by casting and curing a Sylgard 184 mixture of elastomer (Dow corning, USA) and curing agent (10:1, w/w) at 95□. The molded PDMS piece and glass coverslip were placed in a plasma etcher for 1 min (50 W). Then, the molded PDMS piece was placed immediately on the coverslip to form micro-structures.

After the bonding step, the Teflon-AF solution (DuPont™, USA) diluted in FC-40 (3%, v/v, Sigma, USA), was introduced into the culture chambers and micro-bioreactor was placed in a vacuum chamber (0.02 atm) for 10 min to promote adherence to the PDMS surface. Then, the device was incubated at 80°C, 1 hour in a dry oven to evaporate the fluoro-inert solvent to form a Teflon-AF layer. When the solutions had vaporized fully, the PDMS chambers were coated successfully with Teflon.

#### Strains and pre-cultivation

Four species of microalgae from the Key laboratory of experimental marine biology, the Institute of Oceanology, Chinese Academy of Sciences, were grown in 100 ml of natural seawater with f/2 medium in acid washed 250 ml Erlenmeyer flasks. Their main characteristics were shown in Table 1. The natural seawater was obtained from the Bay of Baishi, Dalian, filtered by GF/F Whatman and sterilized by autoclaving. The cultures were incubated at 20±1 °C and under light at 100 μmol photons m^-2^ s^-1^ with a 12 h light: 12 h dark cycle. After 7-9d until the cells entered the stationary phase, a second inoculation of the suspension diluted the cells to 0.5 × 10^7^ cells ml^-1^ in order to maintain the cells in a healthy state for further cultivation.

**Table 1.** Main characteristics of the four microalgae used in the study.

| Microalgal species | Mean length/$\mu\text{m}$ | Mean width or diatom/$\mu\text{m}$ | Mean volume /$\mu\text{m}^3$ |
| --- | --- | --- | --- |
| i. *Dicrateria zhanjiangensis*<br>Hu. Var. sp. | - | 4.81 | 58.24 |
| ii. *Chlorrella*. Sp. | - | 2.86 | 13.43 |
| iii. *Platymonas helgolandica*<br>var. *tsingtaoensis* | 12.50 | 7.38 | 360.07 |
| iiii. *Platymonas subcordiformis* | 10.71 | 6.94 | 269.95 |

#### Characterizing of On-chip BODIPY-Staining

A stock solution of BODIPY 493/503 (4,4-difluoro-4-bora-3a,4a-diaza-sindacene, Molecular Probes, Inc. Eugene, OR, USA, excitation wavelength 480 nm, emission maximum 515 nm) was prepared at 10μg/ml in 10% dimethyl sulfoxide (DMSO) for use as a fluorescent lipid stain. After 15 min incubation in the dark, imaging experiments were conducted using Olympus IX-71 fluorescence microscope (Olympus, Japan) equipped with DP73 CCD camera.

#### On-chip Batch cultures

Batch cultures of Chlorella sp. (chl-1, KLEMB, IOCAS) were grown in 8 units-microfluidic bioreactor. The seed cultures were in nitrate-free f/2 medium to a cell density of 0.5 × 10^7^ cells ml^-1^. A concentrated sodium nitrate stock solution was used to adjust the columns to initial nitrate concentrations that ranged from 0.22 mM to 17.6 mM. All fluids were loaded into the microfluidic device using syringes (Hamiltion, Nevada, USA), and the flow rates of 5μL min^-1^ were controlled using a syringe pump (Harvard, USA). The Nitrate solutions were infused over 30 min to ensure that a consistent nutrient condition in culture chambers with that introduced from mcirofluidic nutrient supply base. And for each nitrate concentration eight replicate cultures were conducted in parallel. The device was then kept in a sealed transparent PMMA culture container with a controlled temperature of 25±0.5°C. 10mL natural seawater with f/2 medium was stored in the container to maintain a 100% relative humidity environment. Natural air was slowly passed the container through a tube. The light illumination was 80 μmol photon m^-2^s^-1^. When observation, the culture container was placed on the motorized stage of an inverted fluorescence microscope (OLYMPUS IX-71). The experimental information were collected by a CCD camera and quantified by a connected computer coupled with Image-Pro Plus software (version 6.0 for Windows XP; Media Cybernetics).

#### On-chip continuous cultures

Continuous cultures of *Chlorella sp*. (chl-1, KLEMB, IOCAS) were carried out similar to the batch cultures, except nitrate in concentration of 0.88 mM was continuously added to the mcirofluidic bioreactor array. For each unit, by flowing seawater (with nitrate-free f/2 medium) and nitrate solution (with nitrate-free f/2 medium) through each inlet, the 8-outlet nutrient supply base produces 0.00, 0.13, 0.25, 0.38, 0.50, 0.63, 0.75 and 0.88 mM concentrations of nitrate solution into downstream nutrient supply channels and then continuously feeds microalgae in chambers. And for each nitrate concentration eight replicate cultures were conducted in parallel.

#### Chemostat cultures

Exactly 2000 mL of *Chlorella sp*. (chl-1, KLEMB, IOCAS) were grown in a closed column bioreactor with an inner diameter of 10 cm. The seed cultures were in nitrate-free f/2 medium to a cell density of 0.5 × 10^7^ cells ml^-1^. A silicone tube was connected to the bottom of the column to form a U-shaped overflow tube. Air was slowly bubbled into the bottom of the culture. A peristaltic pump with a ten-roller pump head was used for continuous mode by feeding f/2 medium (0.18 mM sodium nitrate). Cultivations with different dilution rate of 1.0 d^-1^ was carried out successively. Biomass was determined by measuring of the optical density and biomass dry weight. The lipid content was measured by total lipid extraction method.

## ACKNOWLEDGMENT

Dr. Xingcai Zhang and Dr. Ming Guo acknowledge the support from Harvard/MIT. All others acknowledge the support from National Natural Science Foundation of China (No. 41476085, No. 81471807), Dalian Science &Technology Bureau (Dalian Science and Technology Innovation Fund 2019J12SN55), General Program of Liaoning Science &Technology Department (2021-MS-345), Major Scientific Project of Interscholastic Collaboration of Universities of Liaoning (No. JYT-dldxjc-202001).

## Supporting Information

**Figure s1.**
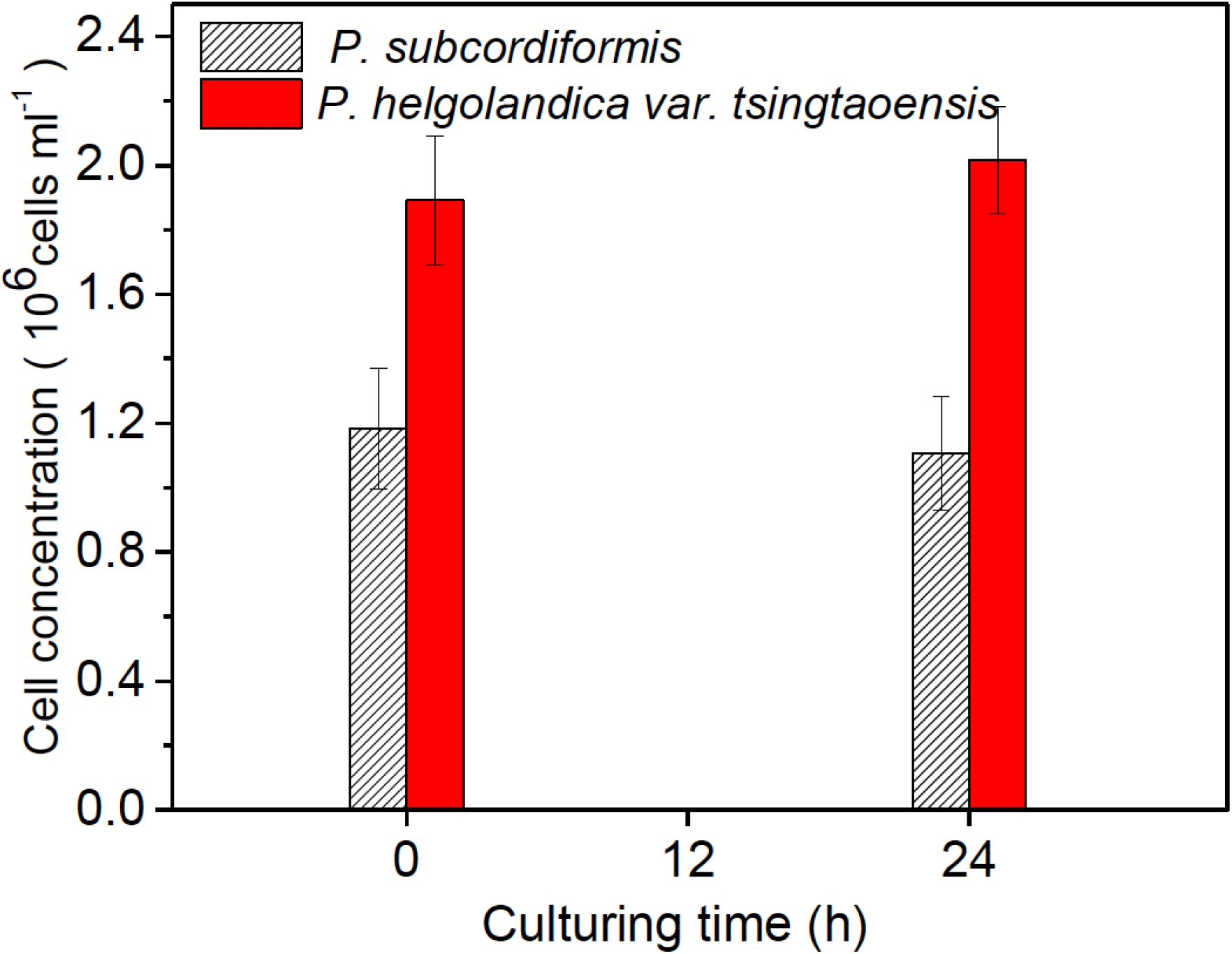
Confining ability assay for microalgae on chip.

**Table s1.** Comparison of growth and lipid accumulation in chemostat and batch culture of *Chlorella sp*. (chl-1, KLEMB, IOCAS)

| Culture mode | Dilution rate | Biomass productivity<br>(mg l<sup>-1</sup> d<sup>-1</sup>) | Lipid content<br>(%DW) | Lipid productivity<br>(mg l<sup>-1</sup> d<sup>-1</sup>) |
| --- | --- | --- | --- | --- |
| **Chemostat** | 1.0 | 353.82 | 18.34% | 64.89 |
| **Batch** | - | 146.57 | 32.10% | 47.04 |

